# Automated segmentation of fetal intracranial volume in 3D ultrasound using deep learning: identifying sex differences in prenatal brain development

**DOI:** 10.1101/2022.12.19.521094

**Authors:** Sonja MC de Zwarte, Jalmar Teeuw, Jiaojiao He, Mireille N Bekker, Ruud JG van Sloun, Hilleke E Hulshoff Pol

**Affiliations:** Department of Psychiatry, UMC Brain Center, University Medical Center Utrecht, Utrecht University, Utrecht, The Netherlands; Department of Obstetrics, University Medical Center Utrecht, Utrecht, The Netherlands; Lab of Biomedical Diagnostics, Department of Electrical Engineering, Eindhoven University of Technology, Eindhoven, The Netherlands; Department of Experimental Psychology, Helmholtz Institute, Utrecht University, Utrecht, The Netherlands

**Author notes:** Correspondence, Dr. Sonja de Zwarte.

**Keywords:** convolutional neural network, 3D ultrasound, automated segmentation, fetal brain development, intracranial volume, sex differences

## Abstract

The human brain undergoes major developmental changes during pregnancy. Three-dimensional (3D) ultrasound images allow for the opportunity to investigate typical prenatal brain development on a large scale. Here, we developed a convolutional neural network (CNN) model for automated segmentation of fetal intracranial volume (ICV) from 3D ultrasound, and we applied the trained model in a large independent sample (N = 9795 ultrasounds; N=1763 participants) from the YOUth Baby and Child cohort measured at 20- and 30-week of gestational age to investigate sex differences in fetal ICV and ICV growth. 168 3D ultrasound images with ICV annotations were included to develop a 3D CNN model for automated fetal ICV segmentation. A data augmentation strategy provided physical variation and additional data to train the network. K-fold cross-validation and Bayesian optimization were used for network selection and the ensemble-based system combined multiple networks to form the final ensemble network. The final ensemble network produced consistent and high-quality segmentations of ICV. Our trained model successfully predicted ICV and ICV growth in a large independent longitudinal data set. Boys had significantly larger ICV at 20- (B=2.83; *p*=1.4e-13) and 30-weeks of pregnancy (B=12.01; *p*=2.0e-28), and more pronounced ICV growth than girls (t=-4.395; *p*=1.3e-5). Our approach provides us with the opportunity to investigate fetal brain development on a much larger scale and to answer fundamental questions related to prenatal brain development.

## 1. Introduction

Ultrasound imaging is widely used among pregnant women to monitor fetal growth and development due to its low-cost, non-invasive, and real-time characteristics. Currently, three-dimensional (3D) ultrasound examinations have been increaisngly used to evaluate the fetal central nervous system, allowing for whole brain imaging at once, instead of one slice at a time (1). Intracranial volume (ICV), as a global measure of structural fetal brain development, is an important assessment tool in 3D ultrasound imaging studies (2–4). During this period of complex developmental processes, the brain is vulnerable for (gene-by-) environmental factors which can have a permanent effect leading to developmental disorders later in life (5–8). However, whether individual differences in brain structure precede problems later in life and at what point during development (i.e., prenatal or postnatal) these differences arise remains largely unknown.

Indeed, normal fetal brain size and growth can inform functional outcome postnatally (9,10). Recently, a more pronounced fetal brain size and growth in early-to mid-gestation was positively associated with academic attainment in mid-childhood learning capacity in the area of mathematics, writing, reading, and logical thinking (11). Moreover, a faster growth from mid to late pregnancy predicted a lower risk of delayed infant development at 12 months of age (12). In addition, a large and steadily accumulating body of evidence suggests that abnormal prenatal brain development is associated with several psychiatric disorders, such as in psychosis (5). However, despite remarkable progress that has been made in understanding the development of the human fetal brain (e.g., Allen Brain atlas), our knowledge on typical human fetal brain development is still limited. To answer questions related to fetal brain development and outcome later in life, several thousand fetal brain measures are required, preferably including multiple measurements to study growth in utero, also when influences of genes and environmental are studied.

Recent work has demonstrated the remarkable performance of deep learning techniques on automated biometrics measurement and segmentation of brain structures in both 2D and 3D ultrasound images (13–24). Moreover, measurements generated in near real time by deep learning techniques could speed up clinical workflow. Accurate automated image segmentation is also a prerequisite for the quantitative assessment of the fetal brain in large-scale studies.

Here, we built a neural network for automated fetal brain extraction and ICV measurement from 3D ultrasound images acquired at 20 and 30 weeks of gestation in a longitudinal setup. We performed a data augmentation and k-fold cross validation with the Bayesian Optimization strategy. Multiple networks were combined as an ensemble network to achieve optimal automated segmentation of fetal ICV at both 20 and 30 weeks of gestation. In a large independent dataset, the model was applied to investigate whether the model successfully predicted ICV by investigating sex differences in ICV development.

## 2. Methods

### 2.1 Study sample and image acquisition

The 3D ultrasound scans were obtained from the ongoing YOUth Baby and Child cohort study (https://www.uu.nl/en/research/youth-cohort-study) (25). In this cohort, over 2,800 children from the greater Utrecht region in the Netherlands are being followed from the fetal stage to childhood. During pregnancy, ultrasounds were acquired at two time points, i.e., around 20 and 30 weeks of gestation. The sweeps were made by experienced sonographers with a Voluson E10 (GE Healthcare, Zipf, Austria). To acquire a 3D ultrasound image, the probe sends out a sweep with sound waves at different angles and the returning echoes are processed and reconstructed in a multiplanar view of the three two-dimensional (2D) orthogonal planes (26).

### 2.2. Image preprocessing

Of 168 high quality 3D ultrasound images of typical developing fetal brains with corresponding ICV annotation were included for the development of the 3D CNN model to automatically segment fetal ICV (27). Next, the model was applied to a large independent set from the YOUth Baby and Child cohort. On average each participant had 10 ultrasounds sweeps acquisitions per measurement, resulting on over 50,000 ultrasound images. All Kretz ultrasound volumes were automatically converted to NIFTI with SlicerHeart extension for 3D Slicer (28), with a fixed isotropic voxel size of 0.6 mm. A crop and orientation step was implemented to align the fetal brains. After prediction a semi-automated quality control procedure was performed, resulting in successful ICV prediction in N = 1763 participants (age 140 to 174 gestational days at baseline, 200 to 229 gestational days at follow-up; 50.7% boys).

### 2.3 Network pipeline

The 3D ultrasounds with corresponding annotations were loaded. Data were split to train data (N = 116, evenly from both age groups) and test data (N_20weeks_ = 29 and N_30weeks_ = 23). In total, 73 subjects were included with both a 20- and 30-week ultrasound. To prevent bias, we ensured that subjects with two scans were not both part of the train and test data.

#### 2.3.1 Data augmentation

A 3D data augmentation strategy was applied three times to the train data, which increased the number of images as well as annotations by random spatial augmentations (elastic, rotation, scaling, mirroring deformation), color augmentations (contrast, brightness, gamma transformation), and noise augmentations (Gaussian noise addition, gaussian blurring transformation) (Figure 1) (29). In addition, this strategy promotes convergence toward model parameterization that is invariant to several common variations (17).

**Figure 1:**
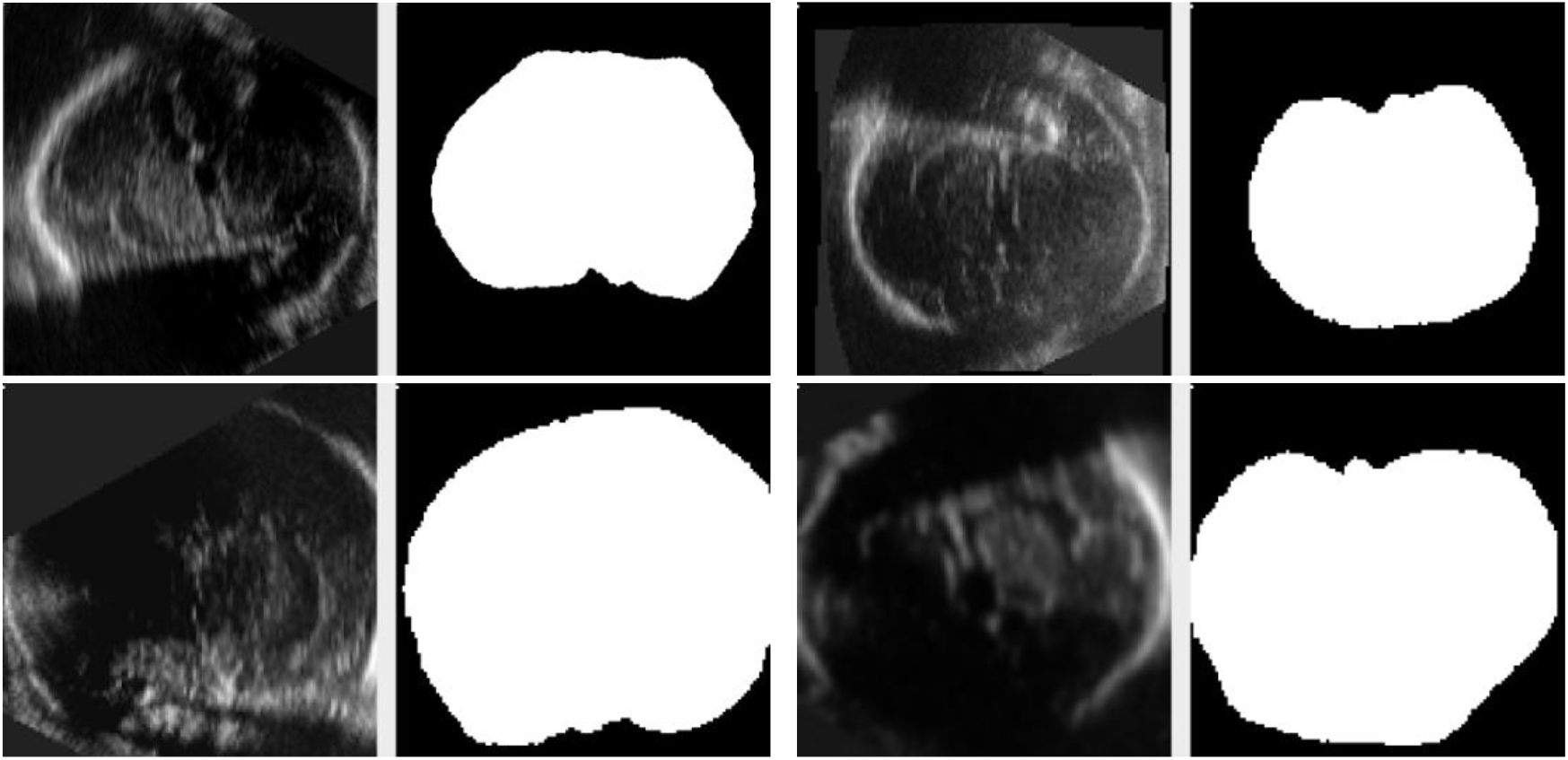
Examples of data augmentation in one participant at 30 weeks of gestation. For each subplot, there is one slice of 3D ultrasound image with corresponding 3D annotation. Top left: the original image and annotation. Top right: the augmented data included mirroring. Bottom left: the augmented data included, e.g., scaling. Bottom right: the augmented data included mirroring, scaling, blurring.

After data augmentation, the train data set included N = 464 3D images. All data, including original data and augmented data, were resized to 128 *×* 128 *×* 128 voxels, and then each color channel of one voxel of 3D images was normalized to numeric values from 0 to 1 while that of 3D annotation was normalized and converted to binary values, i.e., 0 or 1. To balance the train data among the 20-and 30-week groups, the train data set consisted of the same amount of 20- and 30-week data partitioned at an interval of 2 images. The augmented data and nonaugmented data were also distributed at an interval of 4 images.

#### 2.3.2 Convolutional Neural Network architecture

The 3D CNN architecture comprises of encoder and decoder with skip concatenation between them. The architecture contains the convolutional block with batch normalizations and ReLu activations, downsampling/upsampling layers, dropout layers and a softmax layer before output (Figure 2). The network was trained with Adam optimizer, Sparse Categorical Cross-Entropy loss function and two metrics, i.e., voxel-wise accuracy and Dice Similarity Coefficient (DSC).

**Figure 2.**
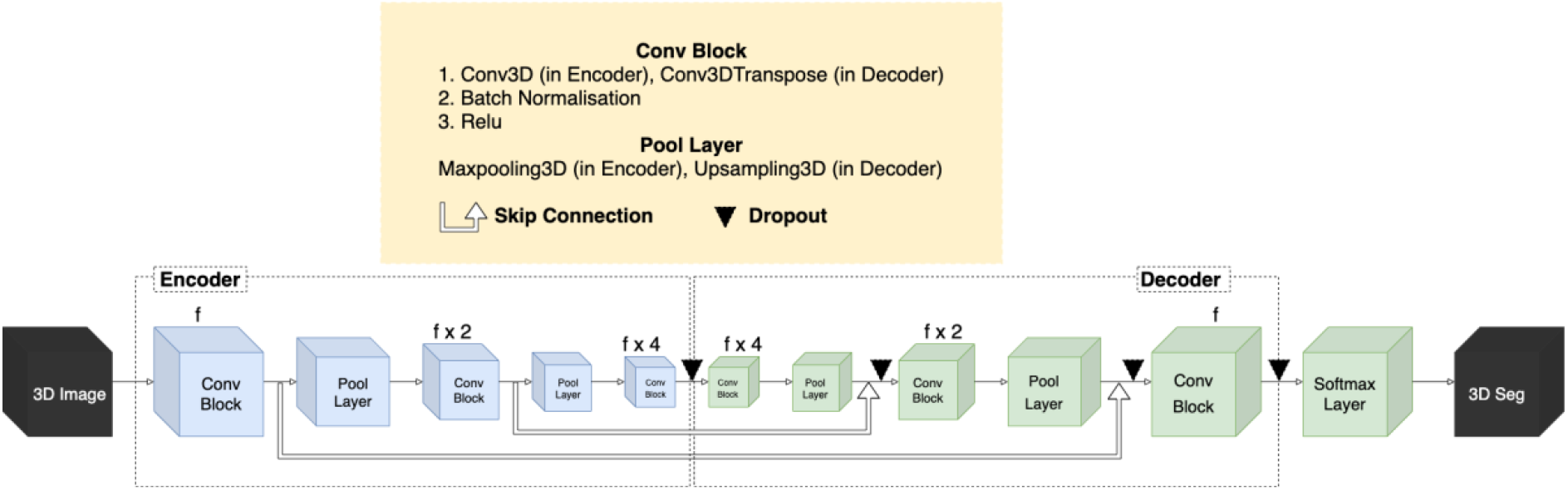
The structure of the 3D convolutional neural network. The Convolutional (Conv) Block starts with one pre-defined filter size (f) and then a multiple of it in the flow from 3D images to 3D segmentations. The kernel size inside the convolution layer is 5 or 3 and the pooling size of the sampling layer is 2.

#### 2.3.3 Network selection

We performed 3-fold cross-validation and Bayesian Optimization based on DSC of validation data to tune network hyperparameters for batch size, filter size and learning rate values. The hyperparameters were set to be optimized for eight iterations by randomly exploring and adjusting parameters based on the previous random points, as well as simultaneously conducting the 3-fold cross-validation scheme. For each fold, voxel-wise accuracy and DSC were calculated during training, and Wilcoxon signed-rank tests were performed to identify the best performing network for each fold with statistical significance. These three networks were combined as an ensemble network by a voting-based technique (i.e., for each voxel the predicted label is assigned by the value with the most votes from sub-networks) (30,31).

#### 2.3.4 Performance assessment

A total of 52 3D images were predicted by the ensemble network and assessed by four metrics (voxel-wise accuracy, DSC, Hausdorff Distance voxel distance (HD_voxel_) and Hausdorff Distance physical distance in millimeter (HD_physical_)). Wilcoxon rank-sum tests determined whether the ensemble network performed differently for the 20- and 30-week age groups. All statistic tests considered *p* < 0.05 as significant. If the value was 0.05 quantile out of one side of the distribution, the outliers were extracted and removed from the test data set. Each group per metric distribution except HD_physical_ was checked for outliers as the relative contour distance under the same voxel volume was more valuable to consider than the absolute distance under different physical volumes.

#### 2.3.5 ICV measurement

ICV *(V)* was calculated by directly counting the voxels of predicted segmentations *(N)*, multiplying the voxel size *(S;* magnitude per voxel):

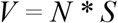

### 2.4 Statistics

All statistical analyses were performed using R (http://www.r-project.org) Linear regression analysis of the intracranial volumes was performed with the nlme package in R at each wave separately (http://CRAN.R-project.org/package=nlme) (32). The models included fixed effects for sex, linear age, and quadratic age, and random intercepts for each subject to account for the repeated measures at each assessment.

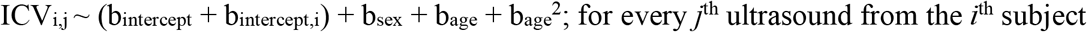

Linear change rates in intracranial volume were computed as the difference between the intracranial volume at 30 weeks follow-up and 20 weeks baseline. The change rates were converted to daily change rates to account for sampling interval between assessments. Daily linear change rates were compared between sexes with a students’ t-test.

## 3 Results

### 3.1 Model performance

#### 3.1.1. Hyperparameter Optimization and Cross-Validation

We divided each type of produced hyperparameters into two groups for batch size and three groups for filter size and learning rate by the constraints commonly used by others since we wanted to narrow the range of hyperparameters to investigate which group performed better and consistently. The 24 networks were trained on three validation folds and were set to different hyperparameters (Figure 3). By comparing the mean of DSC with observation and Wilcoxon signed-rank test, the selected networks outperforming all other networks on each validation fold were No. 5, 13, and 17 (Figure 3, 4). To obtain a powerful and consistent model, we combined the three networks as an ensemble network with a voting scheme to reduce the impact of variation between new datasets and improve the quality of segmentation.

**Figure 3.**
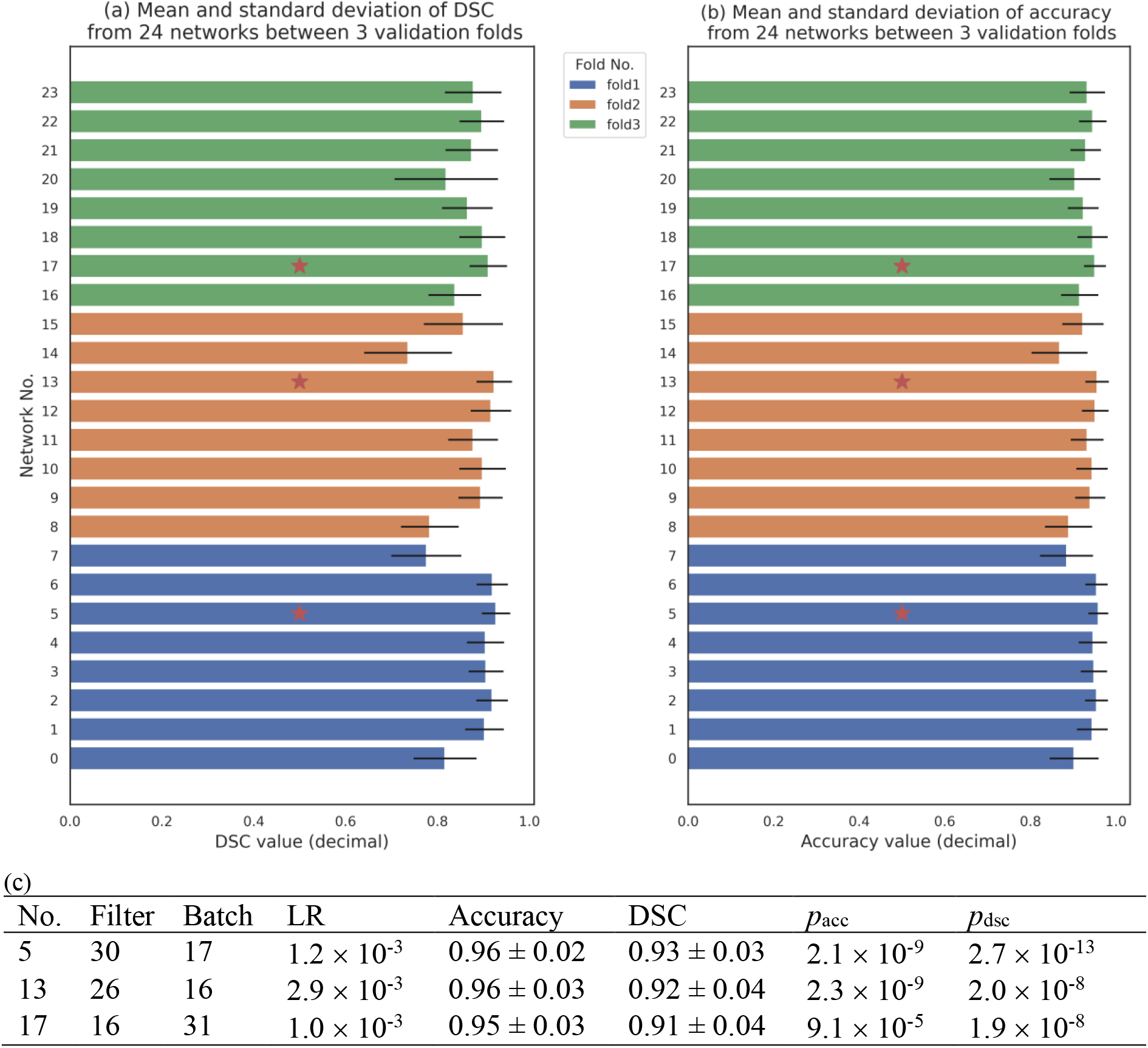
The performance during training of the 24 networks. The mean and standard deviation of (a) dice similarity coefficient (DSC) and (b) voxel-wise accuracy on validation data in each of the 24 networks are depicted by the bars and lines respectively. Eight networks were grouped per cross validation: fold 1 (blue), fold 2 (orange) and fold 3 (green). The red star markers refer to the best performance per metric per fold. (c) The hyperparameters and performance of the best three networks of each fold. The filter size, batch size and learning rate (LR) of networks No. 5, 13 and 17 are in [16, 32), [16, 32) and [0.001, 0.005) respectively. The p-values of voxel-wise accuracy (*p*_acc_), and DSC) (*p*_dsc_), indicate the maximum values for the comparison of three networks to other networks in the same cross-validation fold.

**Figure 4.**
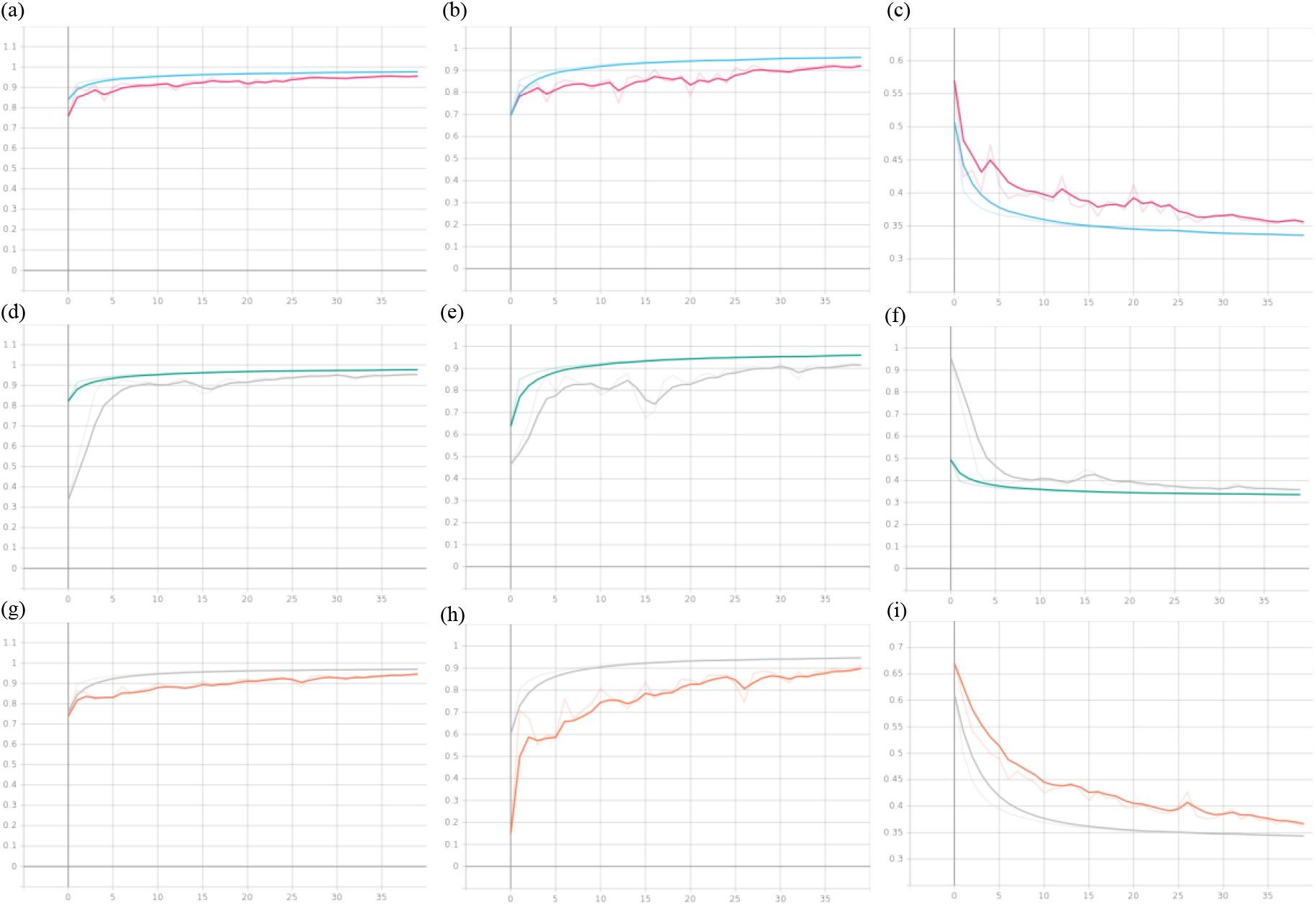
The performance of the best three networks; No.5 (a, b, c), No.13 (d, e, f) and No.17 (g, h, i) over 40 epochs. The left column depicts the voxel-wise accuracy dynamic, the middle column for the Dice Similarity Coefficient (DSC) dynamic and the right column for the Sparse Categorical Cross-Entropy loss dynamic measuring the similarity/difference between annotation and prediction data during training. In each subplot, the two curves represent the train data (blue, green, gray) and validation data (purple, gray, orange) respectively.

#### 3.1.2 Test Data

ICV was predicted in 52 3D ultrasounds as part of the test data set and the findings were assessed by computing the voxel-wise accuracy, DSC, HD_voxel_ and HD_physical_ metrics for the 20- and 30-week age groups. Voxel-wise accuracy and DSC indicate highly correct classification, and HD_voxel_ and HD_physical_ show a small distance of contour for all test data and each gestational age group (Figure 5). There was no significant difference between the 20- and 30-week test data based on DSC (*p*=0.33) and HD_voxel_ (*p*=0.49), while the voxel-wise accuracy did show a significant difference in performance between the 20- and 30-week age group (*p*=0.03). The reason for the significant group effect in voxel-wise accuracy might be that the background accounts for a much larger proportion for the earlier gestational ages and voxel-wise accuracy considers background as part of the calculation, while DSC and HD do not take the background into account.

**Figure 5.**
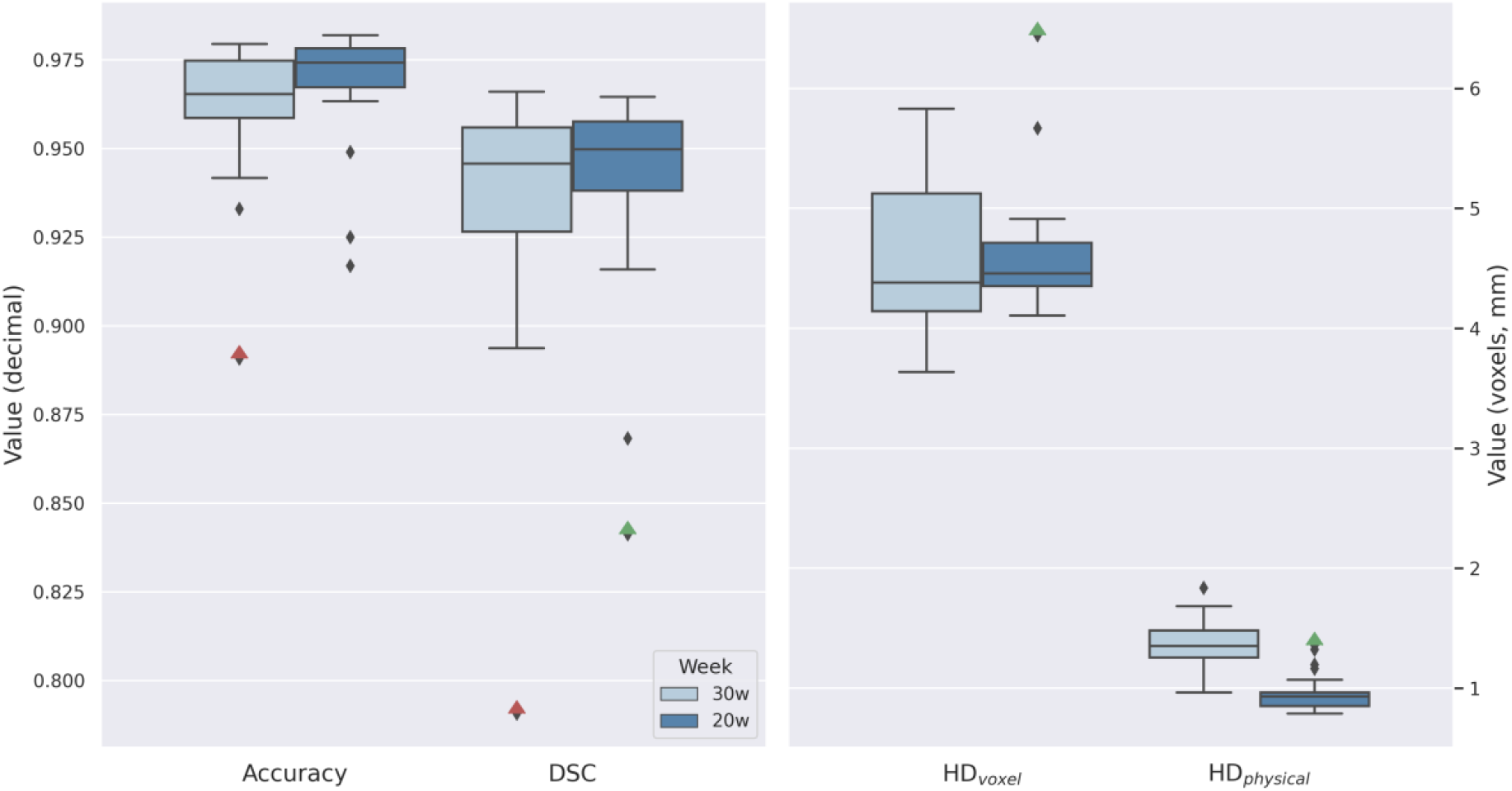
The distribution of 20-week (dark blue) and 30-week (light blue) test data for four metrics, i.e., voxel-wise accuracy and dice similarity coefficient (DSC), and the right y-axis indicate the scale of Hausdorff Distance in voxels (HD_voxel_) and physical HD in millimeters (HD_physical_). The red and green triangles represent for two recurring outliers.

### 3.2 Sex differences in ICV development

Our model successfully predicted ICV measures in N = 9795 ultrasound images from 1763 fetuses (N_20weeks_ = 1297, age range = 140-174 days, 53.0% boys; N_30weeks_ = 1315, age range = 200-229 days, 48.3% boys), who were not part of training, validating, or testing of the model. This was ~20% of the available ultrasound scans. Most unsuccessful predictions were due to non-brain ultrasound, low-quality ultrasound (e.g., due to motion of fetus or high maternal BMI), or incomplete coverage of the entire fetal brain.

Boys had significantly larger ICV at 20- (B=2.83; *p*=1.4e-13) and 30-weeks of pregnancy (B=12.01; *p*=2.0e-28) (Table 1, Figure 6, 7). Moreover, boys showed more pronounced ICV growth than girls (t=-8.394; *p*<2.2e-16) with an annualized growth rate of an average 3.15ml vs 3.10ml per day (Table 1, Figure 8).

**Figure 6.**
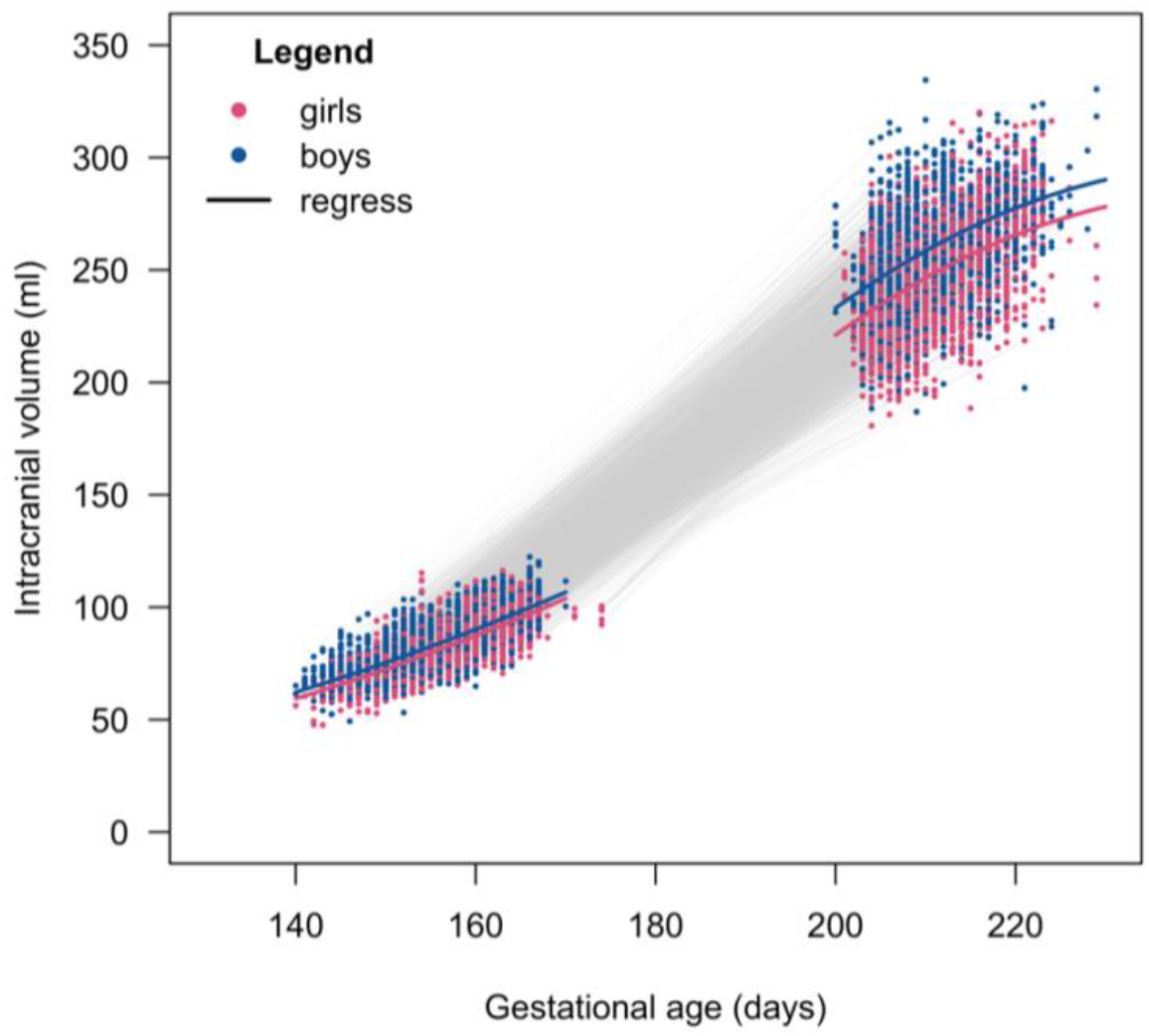
Growth curve for fetal ICV of boys and girls. Models included a quadratic age effect and sex effect.

**Figure 7.**
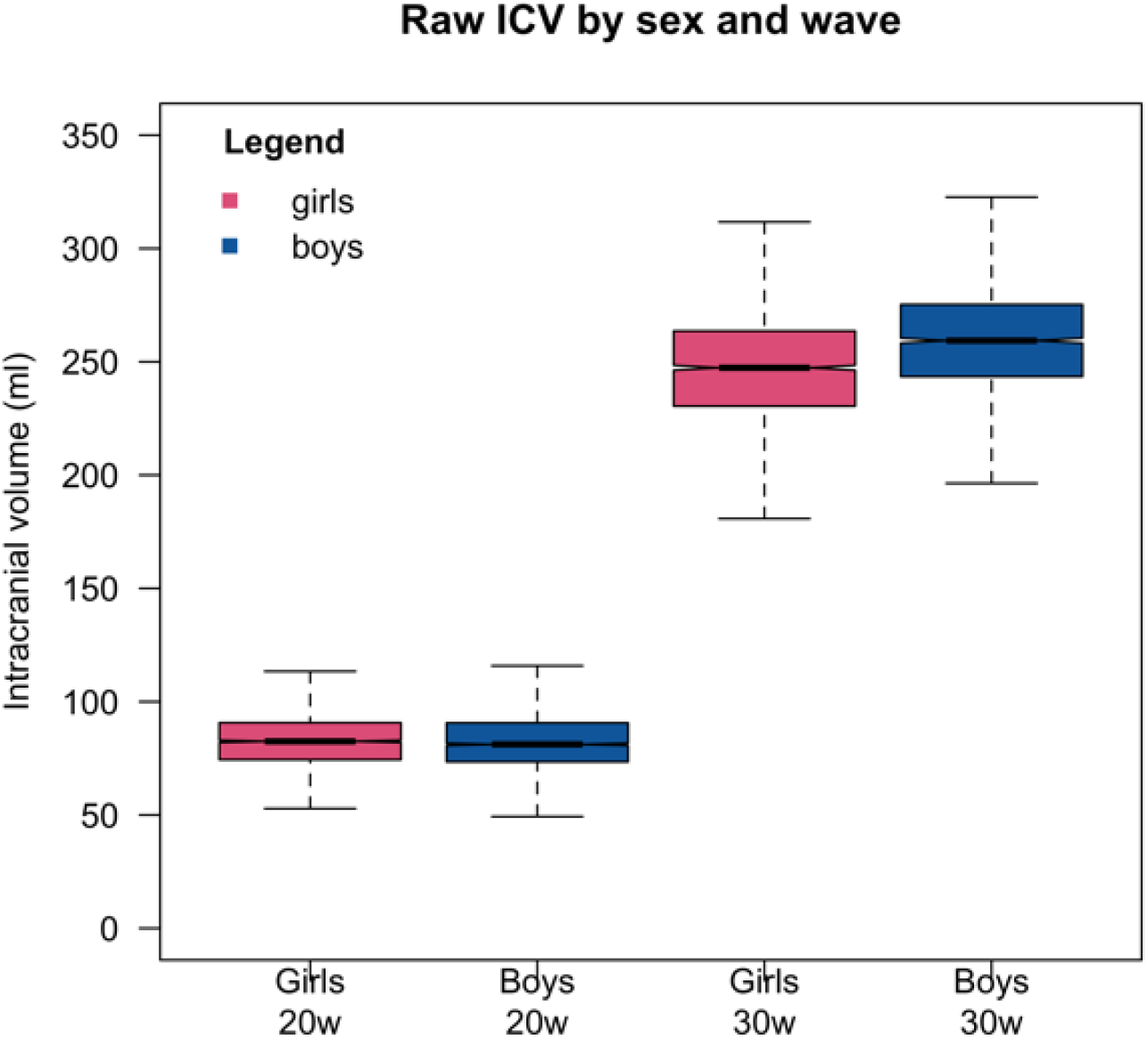
Sex effect for fetal ICV of boys and girls at baseline 20 weeks and follow-up 30 weeks. Sex effects were significant at baseline 20 weeks and follow-up 30 weeks after correcting for linear and quadratic age effects.

**Figure 8.**
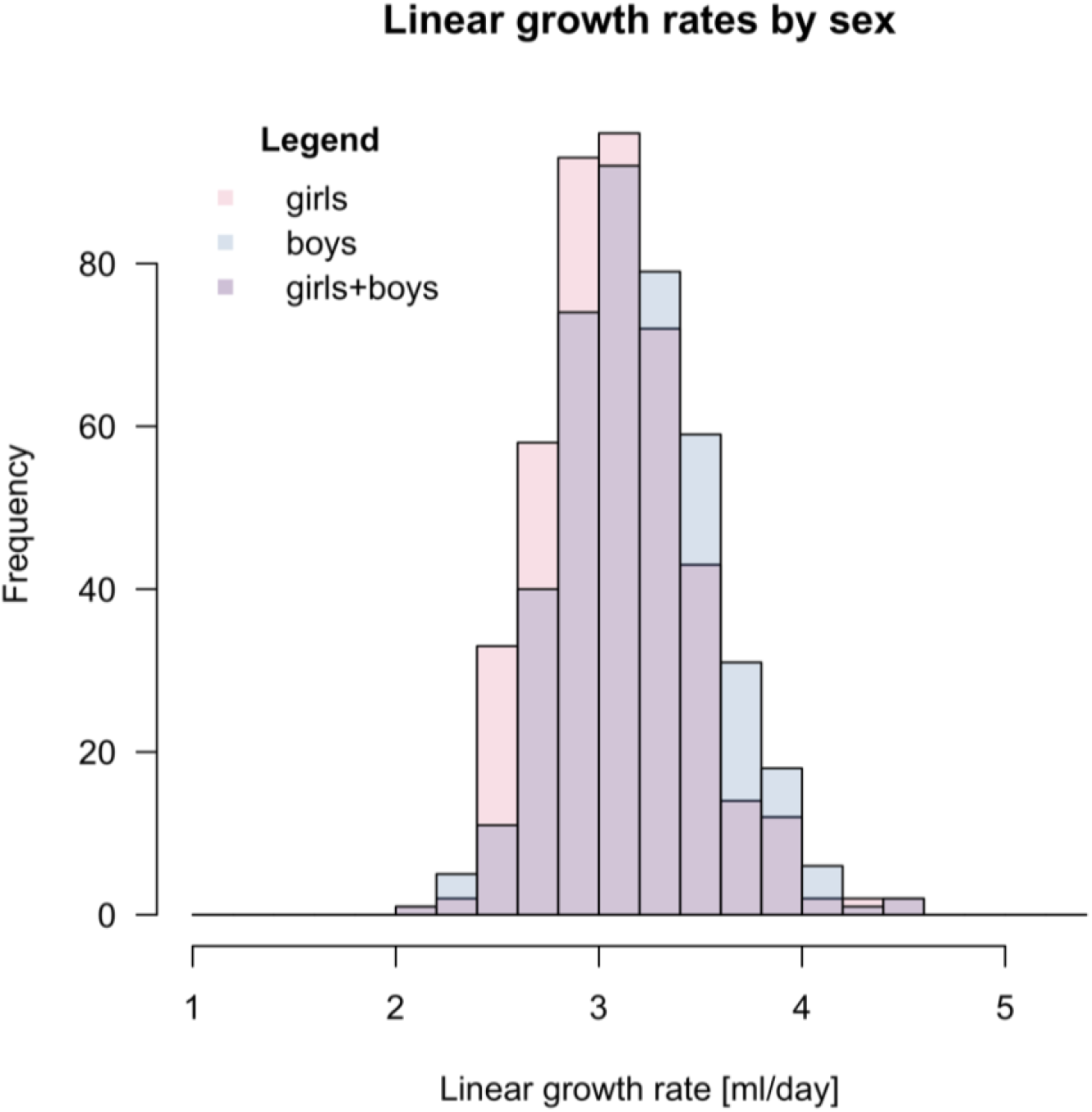
Daily linear growth rates for boys and girls between baseline 20 weeks and follow-up 30 weeks assessment.

**Table 1.** Regression coefficients of sex and linear- and quadratic age effects on fetal ICV at baseline 20 weeks and follow-up 30 weeks. The B-coefficients and standard error (SE) are reported together with nominal *p*-values. A positive sex effect indicates larger ICV volumes for boys compared to girls. Longitudinal growth rates were computed by the difference in ICV volume at 30 weeks follow-up subtracted by the ICV volume at 20 weeks baseline assessment.

|  | **Baseline**<br>**20 weeks** | **Follow-up**<br>**30 weeks** | **Longitudinal**<br>**growth rate** |
| --- | --- | --- | --- |
| Intercept | 55.60±105.03 | -1628.35±713.10 | N/A |
| Sex | 2.83±0.38<br>**$p=1.4\text{e-}13$** | 12.01±1.06<br>**$p=2.0\text{e-}28$** | $t = -4.395$; $df = 844$<br>**$p=1.3\text{e-}05$** |
| Age | -1.17±1.36<br>$p=0.388$ | 15.63±6.71<br>**$p=0.020$** | N/A |
| Age<sup>2</sup> | 0.01±0.00<br>$p=0.051$ | -0.03±0.02<br>**$p=0.043$** | N/A |

## 4. Discussion

In this study, we report on a deep learning method using a convolutional neural network for automated segmentation and measurement of fetal ICV in 3D ultrasound images. Our automated tool was able to produce good-quality segmentations of ICV, which seemed to overcome challenges such as the fetal pose and differences in gestational age. In addition, we were able to identify significant sex differences in fetal ICV and ICV growth with our developed model in a large independent sample. We show that boys have a larger ICV at both 20 and 30 weeks of gestational age and also have more pronounced ICV growth.

To the best of our knowledge, we are the first longitudinal ultrasound study to show sex differences in fetal ICV development in almost 10,000 ultrasound images from 1763 fetuses. Our findings are in line with previous ultrasound studies investigating in-utero head circumference (as a 2D proxy for ICV), where sex differences were reported as early as 15 weeks of gestation (33–37). Our findings also support and extend the well-known finding that the median head circumference of males at birth is larger than those of females (38). Sexual dimorphisms are seen postnatally in behavior, language development and neurodevelopmental disorders (e.g., autism spectrum disorder and attention deficit hyperactivity disorder) (39). Boys had a larger ICV when compared to girls in-utero already at 20 weeks of gestation confirming that sex differences occur already early during development. Moreover, boys when compared to girls had a more pronounced growth of ICV between 20 and 30 weeks of gestation. These findings suggest that already around 20 weeks of gestation the brains of boys and girls develop differently, and in the period between 20 and 30 weeks of pregnancy this differential growth becomes more pronounced. Our findings further confirm that these sex differences might have a very early and prenatal origin.

Human brain structure is known to change throughout the lifespan and that it is under the influence of both genes and environment. Recently, we identified common genetic variants that affect rates of brain growth or atrophy in what is, to our knowledge, the first genome-wide association meta-analysis of changes in brain morphology across the lifespan in 15,640 individuals (40). In that study we demonstrated global genetic overlap with depression, schizophrenia, cognitive functioning, insomnia, height, body mass index and smoking. Interestingly, gene set findings implicated early brain development in the rates of brain changes of multiple brain structures. Environmental influences, such as prenatal famine exposure has also been associated with decreased intracranial volume (8). Identifying variants involved in structural brain changes may help to determine biological pathways underlying optimal and dysfunctional brain development and the differential influences on males and females. Deep learning network analysis in longitudinal 3D ultrasound of fetal brain allows to study the impact of prenatal brain development on postnatal outcome.

Previous deep learning to annotate the fetal brain included varying approaches. A very deep convolutional network for large-scale image recognition (VGG) network was developed and exploited for fetal brain detection and measurement of head circumference and biparietal diameter with an ellipse model fitted to the segmentation respectively (14,15). In addition to brain-extraction, automatic zonal segmentation was also performed with a convolutional network for biomedical image segmentation (U-Net) (16,17). A multi-task fully convolutional network technique was leveraged to the measurement of abdominal circumference, and a 3D fetal brain model was aligned to a canonical reference space (18,19). A 3D convolutional neural network (CNN) approach worked well on accurately and reliably extracting the fetal brain regardless of the large data variations in the acquisition and the pose variation of the subject, the scale, and even partial feature-obstruction in the 3D ultrasound images (13,20,21). The performance of our ensemble network is in line with previous work by Moser et al (13), who proposed a similar CNN for fetal brain segmentation from 3D ultrasound. Indeed, our findings strengthen the potential of CNN for fetal brain segmentation purposes. Moreover, it is important to note that we developed our tool with a relatively small dataset and with differential gestational age.

We showed great performance in our high-quality annotated test data; however, when applying the model to the large cohort data with many raw ultrasound images per person with varying quality and regions of interest, we also encountered some issues. The YOUth Baby and Child cohort data has a specific study design with two repeated measures between roughly 19-23 and 29-33 weeks of gestational age, instead of a continuous distribution of ages. The brain develops rapidly between these two timepoint, with ICV expanding three-fold within these 10 weeks as we confirmed, and with dramatically increased gyrification (41). It seems that our model had the tendency to over-annotate the smaller 20-week brains and under-annotate the larger 30 week brains. As a next step, we will investigate whether we can optimize our model to better account for our longitudinal study design. In addition, in this study we took a classical approach by applying a semi-automated crop and orientation procedure to align the brains. However, our findings conformed that a fully automated alignment step as proposed in Moser et al (20) is preferred and will therefore be included in the next model. Moreover, next steps will also include adding segmentation confidence through a Bayesian Monte Carlo dropout approach (17) and the use high contrast labeled fetal MRI to extend the pipeline for annotations of cortical and subcortical brain structure in ultrasounds (42).

In summary, we presented an automated tool for extracting fetal brain structure from 3D ultrasound, which predicted good-quality segmentation of fetal ICV. Our automated tool performed accurately for 3D ultrasound images acquired between 19-33 weeks of gestational age and provided reliable ICV results. The automated tool provided us with the opportunity to investigate fetal brain structure on a much larger scale, and we were able to show significant sex differences in early brain development. In the future, we would extend our network by predicting multiple classifications for substructures inside the fetal brain or by multi-task learning techniques to explore more biological problems. In a clinical setting, such automated tools might ultimately aid in the screening for potential neurodevelopmental malformations.

## Acknowledgments

YOUth is funded through the Gravitation program of the Dutch Ministry of Education, Culture, and Science and the Netherlands Organization for Scientific Research (NWO grant number 024.001.003). YOUth is part of (and partly funded by) the research theme Dynamics of YOUth of Utrecht University and of the UMC Utrecht Brain Center.

